# No association of mitochondrial DNA levels in trophectodermal cells with the developmental competence of the blastocyst and pregnancy outcomes

**DOI:** 10.1101/629956

**Authors:** G Ritu, Geetha Veerasigamani, Mohammed C. Ashraf, Sankalp Singh, Saniya Laheri, Deepak Modi

**Affiliations:** Department of Medical Genetic, Craft Hospital and research Center, Chandappura, Kodungallur, Pin – 680664, Thrissur District, Kerala. India; Department of Reproductive Medicine, Craft Hospital and Research Center, Chandappura, Kodungallur, Pin-680664, Thrissur District, Kerala. India; Molecular and Cellular Biology Laboratory, National Institute for Research in Reproductive Health (ICMR), J. M. Street, Parel, Mumbai – 400012, India

**Author notes:** Correspondence address. E mail.

**Keywords:** Mitochondrial DNA, Next Generation Sequencing, blastocyst, implantation, ploidy, maternal age, morphology

## Abstract

**Study question:** Can mitochondrial DNA (mtDNA) levels in trophectodermal cells of the blastocyst predict the blastocyst quality, ploidy status, implantation rate and clinical outcomes?

**Summary answer:** mtDNA levels in trophectodermal cells of the blastocyst do not associate with the blastocyst quality, ploidy status, implantation potential and clinical outcomes, but can differentiate between aneuploid and euploid blastocysts.

**What we already know:** mtDNA levels in the trophectodermal cells have been suggested to be associated with blastocyst morphology, ploidy and implantation rates, and has been proposed as biomarker to access blastocyst quality and predict clinical outcomes. However, discrepancies exist if mtDNA levels could serve as a marker for the same.

**Study design and duration:** Retrospective analysis of mtDNA levels in trophectodermal cells obtained from blastocysts undergoing preimplantation genetic testing for aneuploidy (PGT-A) at Craft Hospital & Research Center, Kerala from January 2016-July 2017.

**Participants/materials and methods:** Study included data from 287 blastocyst from (61) couples who underwent PGT-A using next generation sequencing (NGS). Levels of mtDNA in trophectodermal cells of the blastocyst were estimated by the NGS. Comparison of mtDNA levels with maternal age, blastocyst morphology, ploidy status, implantation rates, miscarriage rates and live birth rate was done.

**Main results:** The levels of mtDNA in the trophectoderm of the blastocyst did not correlate with maternal age. There was no significant difference in the mtDNA levels between grade 1 and grade 2 blastocyst. Euploid blastocyst had significantly lower amounts of mtDNA levels in trophectodermal cells of the blastocyst were compared to aneuploid blastocyst. No significant differences were seen between mtDNA levels and implanting and non-implanting blastocysts or those resulted into miscarriage or live birth.

**Limitations:** The study is limited by a small sample size and hence type II error cannot be ruled out.

**Wider Implications:** The study does not support the potential use of mtDNA levels in the trophectodermal cells as biomarker for blastocyst quality and predicting clinical outcomes needs.

**Study funding/competing interest(s):** There is no external funding for the study. There is no conflict of interest.

## Introduction

Assisted reproduction (ART) has witnessed significant progress in the last three decades and has benefitted many infertile couples. Despite, the advancement in ART, the take-home baby rates are still low (Sadeghi et al., 2012). Various factors which contribute to success rate of ART include maternal and paternal age, gamete quality, endometrial receptivity and most importantly embryo quality (Colaco and Sakkas, 2018; Miao et al., 2009; Oron et al., 2014; Revel, 2012). The current embryo selection methods rely on assessment of embryo morphology with a subjective grading criteria or a real time monitoring of the embryonic development and assessment of multiple quantitative endpoints. However, none of these have been of great aid in improving pregnancy rates (Bromer and Seli, 2008; Capalbo et.al., 2014; Minasia et.al., 2016). The advent of single cell genetic analysis, has made it apparent that a large number of embryos developed *in vitro* are chromosomally aneuploid (Demko et.al., 2016). This has prompted the evaluation of chromosomal complement of the embryos by preimplantation genetic testing for aneuploidy (PGT-A). While the introduction of PGT-A has led to reduction in miscarriage, the improvements in live birth rate following PGT-A remains relatively low and as many as 50% of embryos diagnosed as “euploid” by PGT-A do not implant (Capalbo et al., 2015; Chang et.al., 2016). Thus, genomic aneuploidy may not be a sole factor responsible for failure of implantation and hence there is a need to identify embryo selection markers for improving pregnancy rates.

Recently, levels of mitochondrial DNA (mtDNA) has emerged as possible marker for embryo selection (Cecchino et al., 2019, Humaidan et al., 2018, Kim & Seli, 2019). Fragouli and group were the first to report that high levels of mtDNA in embryos derived from older mothers or embryos with aneuploid and embryos with higher levels of mtDNA rarely implant (Fragouli et.al., 2015). These initial findings were subsequently confirmed in a large blinded retrospective study by same group. Large cohort studies have suggested, the quantity of mtDNA as a new biomarker to assist embryo selection (Fragouli et.al., 2017; Ravichandran et.al., 2017). The above studies were confirmed by two more groups independently (de Los Santos et al., 2018; Diez-Juan et al., 2015). However, three additional studies have failed to show any such correlation. Treff and group reported no association between mtDNA levels in implanted and non-implanted embryos (Treff et. al., 2017). In another study the authors argued that the differences in mtDNA content could arise out of differences in genomic DNA content, and hence a normalization factor was used, which led to the conclusion that the levels of mtDNA are near identical between blastocysts stratified by ploidy, maternal age, or implantation potential (Victor et.al., 2017). Another study by Qui and group failed to establish any correlation between mtDNA, embryo quality and clinical outcomes (Qui et al., 2018). Thus, the levels of mtDNA in predicting the embryo quality, implantation potential of embryos and pregnancy outcomes remains controversial.

In the current study, we present our experience with use of trophectodermal mtDNA levels to determine blastocyst quality, implantation potential of blastocyst and clinical outcomes that have undergone PGT-A in a clinical setup. Our results reveal no significant association of trophectodermal mtDNA levels with maternal age, implantation potential and clinical outcomes, however mtDNA levels in trophectodermal cells of the blastocyst can differentiate between euploid and aneuploid blastocyst.

## Materials and Methods

This was a retrospective study of next generation sequencing (NGS) workflow data from patients who underwent PGT-A between January 2016 to July 2017 at Craft hospital and Research Center, Kerala, India. The study was approved by the ethical review board of Craft Hospital & Research Center, Kodungallur, Kerala (ethics no: 002/21/3/2019).

### Study group

At the Craft Hospital and Research Center we routinely offer PGT-A to patients with recurrent implantation failure, recurrent pregnancy loss and advanced maternal age (above 35yrs). The patient characteristics are described in Table 1. A total of 61 couples agreed to undergo PGT-A by NGS from January 2016 to July 2017. Written informed consent was obtained from patients/couples prior to the PGT-A.

### Ovarian stimulation, Intracytoplasmic sperm injection (ICSI) and embryo culture

Controlled ovarian stimulation (COS) was done with antagonist protocol using gonadotropins with dosage between 150 – 300IU depending on age and body mass index. Oocytes were aspirated under local anesthesia after 36 h of agonist trigger. Denudation was done after 1h of oocyte retrieval. ICSI was performed as described previously (Velde et al., 1998). After ICSI, fertilization check was done next day, followed by day 3 embryo quality check. Good quality day 3 cleavage stage embryos (embryos with 6-8 cells, equal size of blastomere and cytoplasmic fragmentation less than 10%) were continued to grow till blastocyst stage. Zygotes were cultured in VITROMED culture medium for 5-6 days. Blastocysts quality was graded as described previously (Gardner et al., 2000; Sen et al., 2013). Patients that required oocyte accumulation, the oocytes were frozen within 30 mins after denudation and ICSI done later.

### Embryo/Blastocyst Biopsy and vitrification/freezing down (of blastocyst)

Trophectoderm biopsy was done on day 5 embryos (blastocyst stage) as described previously (Capalbo et al., 2015). Briefly laser assisted hatching was performed using LYKOS Laser (Hamilton Thorne; MA, USA) on day 3 embryo. On day 5, herniating blastocyst were selected, and 5-10 trophectoderm cells were removed by suction followed by laser pulsation. The trophectoderm cells were collected in phosphate buffer saline (PBS) and stored at −80° C until further processing. Vitrification method was used to freeze down the blastocyst. The blastocysts were equilibrated in equilibration solution for 12 to 15 mins, followed by transfer to vitrification solution. This was finally transferred to cryolock containing liquid nitrogen.

### Whole genome amplification and Next generation sequencing for trophectodermal cells

Whole genome amplification (WGA) on each biopsy was performed using the Rubicon PicoPLEX WGA kit (Agilent, CA, USA) as per manufacture’s recommendations. Following WGA, next generation sequencing (NGS) of the trophectodermal biopsies was carried out (Well et al., 2014). For constructing WGA library, Ion Xpress Plus fragment library kit, and Ion Xpress barcode adapters 1–32 kit were performed as per manufacturer’s instructions (Thermo Fisher Scientific). 150 ng of WGA DNA was fragmented to generate 280 base pair fragments using Ion Shear Plus reagent for 4 mins. Purification of fragmented DNA was done with Agencourt AMPure XP reagent beads (Beckman Coulter, CA, USA), followed by barcoded adaptor ligation, nick repair and purification as per manufacturer’s instructions. E-Gel Size Select (Thermo Fisher Scientific) agarose gel was used to select a peak size of 280 base pairs, followed by amplification of DNA with 10 cycles of polymerase chain reaction (PCR) using Platinum PCR SuperMix High Fidelity (Thermo Fisher Scientific). Individual libraries were diluted to 100 pM. On Ion 520 Chip, a pool of 24 samples were loaded. Ion Sphere particles containing amplified DNA were prepared with Ion PI Template OT2 200 Kit v3 (Thermo Fisher Scientific). Template-positive Ion Sphere particles were enriched with the Ion OneTouch ES (Thermo Fisher Scientific). They were later sequenced with Ion 520 Chip and Ion PI Sequencing 200 Kit v3 on the Ion S5 instrument (Thermo Fisher Scientific). Approximately 3 million reads were obtained for each barcoded sample.

### Analysis of mtDNA levels

The ploidy status and mtDNA levels were analyzed for all the blastocysts undergoing PGT-A with NGS was assessed with Ion Reporter Cloud based software 5.3 (Thermo Fisher Scientific, MA, USA). The mitochondrial DNA levels were calculated: / by Ion Reporter Software.

### Embryo transfer

Total of 287 blastocysts were investigated for mtDNA levels in trophectodermal cells, out of which 68 euploid embryos were selected who underwent frozen embryo transfer. The endometrium was prepared for transplantation using hormone replacement protocol. Estradiol valerate was administered from day 2 of cycle, in a dose dependent manner. Serial ultrasound monitoring was done to check endometrium thickness (10mm), after which oral and vaginal progesterone were administered and frozen embryo transfer was done. The vitrified blastocyst in the cryolock were directly placed in transfer solution (TS), followed by washing with sucrose solution at 37°C. Serum beta human chorionic gonadotropin levels were measured using HCG STAT Elecsys assay on Cobas E601 Immunology Analyzer (Roche, Basel, Switzerland) after 2 weeks of embryo transfer to detect biochemical pregnancy. Clinical pregnancy was determined by transvaginal ultrasound was done at 6 weeks to see an intrauterine gestation sac.

### Statistical analysis

Linear regression analysis was done to study the correlation between maternal age at the time of oocyte retrieval and mtDNA from trophectodermal cell of blastocyst. Correlation between mtDNA level and blastocyst morphology, ploidy status and pregnancy outcomes were carried out. One-way ANOVA using Tukey’s all column comparison test was performed using GraphPad Prism, version 5. *p < 0.05* was accepted as statistically significant.

## Results

### No correlation between mtDNA levels in trophectodermal cells of the blastocyst and maternal age at the time of oocyte retrieval

To investigate the correlation between mtDNA levels in the trophectodermal cells of blastocyst and maternal age at the time of oocyte retrieval, mtDNA levels were analyzed from trophectodermal cells of blastocyst generated/collected from 61 women (between the range of 24-46 with an average age of 32.6 ± 4.13). Linear regression analysis did not show any correlation between mtDNA levels in the trophectodermal cells of blastocyst and maternal age at the time of oocyte retrieval (Fig.1).

**Figure 1.**
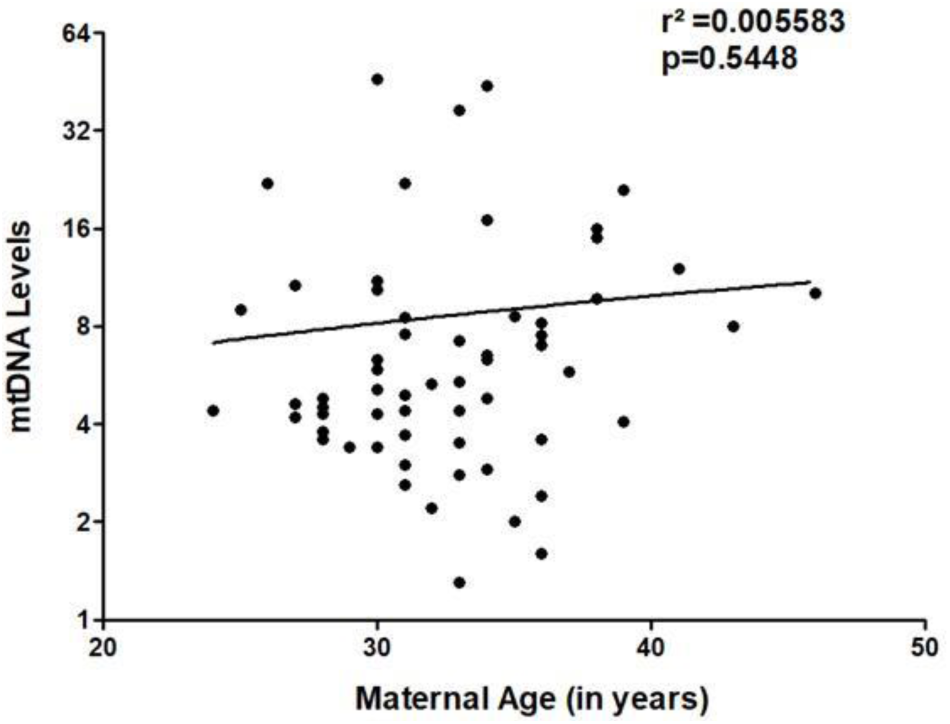
Correlation of mitochondrial DNA (mtDNA) levels in trophectodermal with maternal age at the time of oocyte retrieval. Values on X axis are maternal age in years. Y axis is mtDNA levels in the trophectodermal cells. Each dot represents data form one patient. The data is derived from 61 independent samples. p < 0.05 was accepted as statistically significant.

### mtDNA levels in the trophectodermal cells of the do not correlate with blastocyst morphology

The predominant criteria for selection of blastocyst is morphological grading of the blastocyst. In order to evaluate if there was a correlation between the mtDNA levels in the trophectodermal cells and grade 1 (n= 131) and grade 2 (n= 151) blastocyst were analyzed. PGT-A was not performed on grade 3 blastocyst and thus mtDNA levels are not calculated. The results revealed no difference in the mtDNA levels between grade 1 and grade 2 blastocysts (Fig.2A).

### mtDNA levels in trophectodermal cells is higher in aneuploid blastocysts

To study the correlation between mtDNA levels and ploidy status, the blastocysts were classified into euploid and aneuploid. Out of 287 blastocysts analyzed, 170 were euploid and 117 were aneuploid. Among 117 aneuploid blastocyst, 41 were monosomic, 31 were trisomic and 45 were polysomic (aneuploidy of more than 1 chromosome), mtDNA levels were significantly high in aneuploid blastocysts compared to euploid blastocyst (p <0.05, Fig. 2B). The mtDNA levels were significantly higher only in polysomic blastocysts as compared to euploid and monosomic (p< 0.05, Fig 2C)

**Figure 2.**
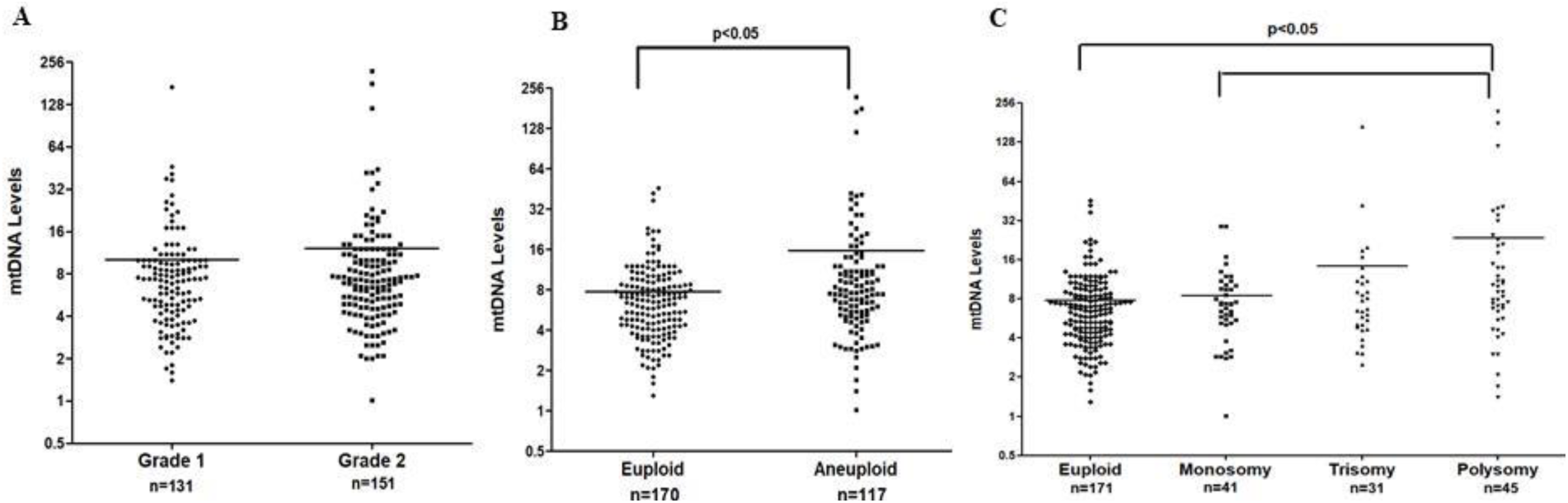
Association of levels of mitochondrial DNA (mtDNA) in trophectodermal cells with blastocyst morphology and ploidy status. (A) Comparison of mtDNA levels in Grade1 and Grade 2 blastocysts. (B) Comparison of mtDNA levels in enploid and aneuploid blastocysts (C) Comparison of mtDNA levels in enploid, monosomic. trisomic and polysomic blastocyst. In all the graphs, values on Y axis is mtDNA levels estimated by next generation sequencing. The numbers (n) of blastocysts are given in each case. Each dot represents data form one patient, p < 0.05 was accepted as statistically significant.

### Levels of mtDNA in trophectodermal cells of blastocyst do not correlate with embryo implantation rate and clinical outcomes

Implantation rate is defined as the number of gestational sacs seen at 6weeks of gestation in ultrasound divided by the total number of embryos transferred. All the couples had atleast one euploid embryo for transfer. Of these 4 couples had double embryo transfer. The remaining 57 couples had single embryo transfer. Out of the 68 blastocyst, 46 blastocysts implanted, whereas 22 blastocysts failed to implant. The overall implantation rate was 67.6%. There was no statistical difference in levels of mtDNA of implanted blastocysts vs non-implanted. The couples who were positive for clinical pregnancy were followed up until end of term. There was no statistical significance difference in mtDNA levels and pregnancy that resulted in miscarriage or live birth (Fig 3).

**Figure 3.**
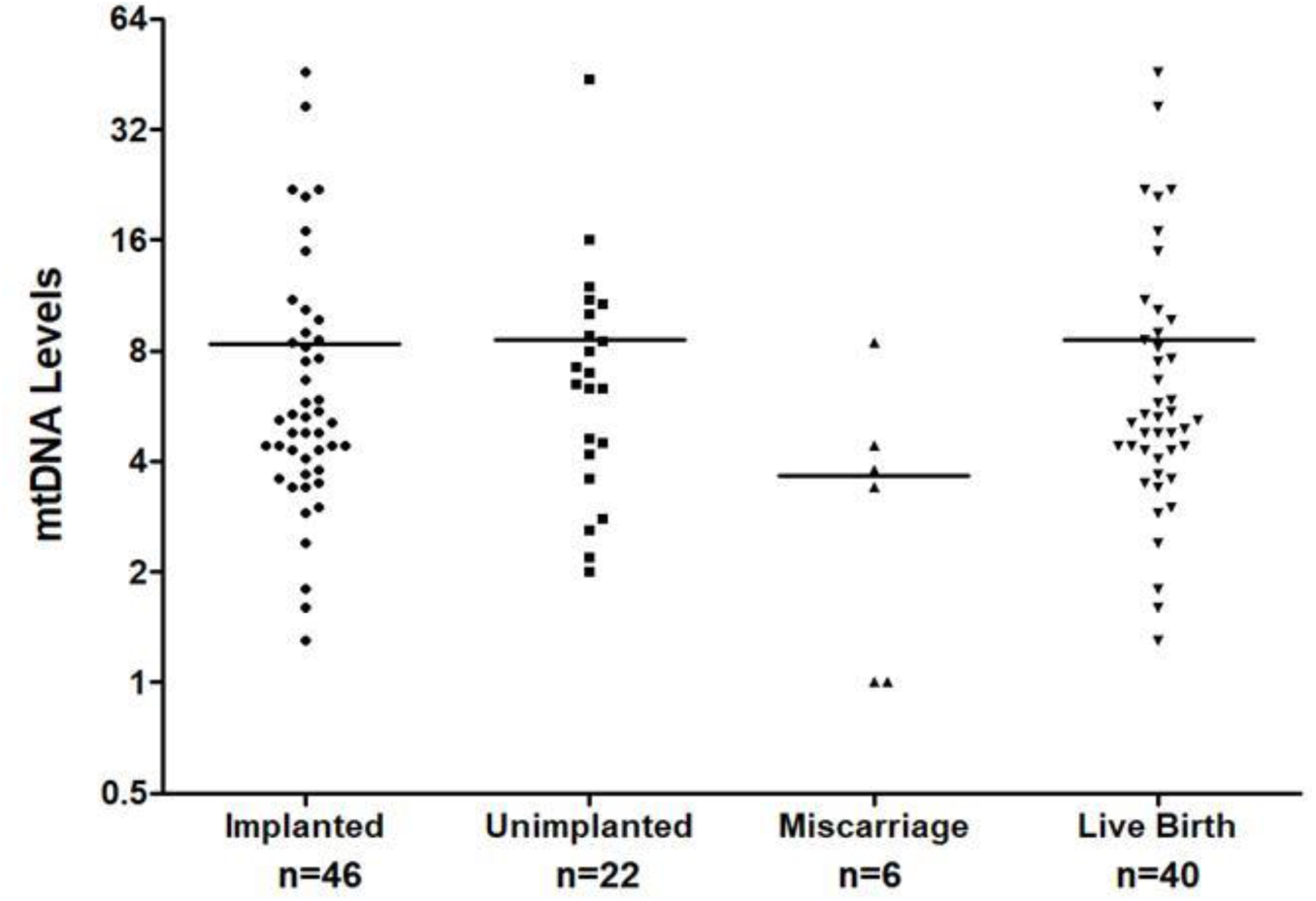
Association of mitochondrial DNA (mtDNA) levels in trophectodermal cells implanted euploid embryos, and clinical outcomes. Values on Y axis is mtDNA levels. The numbers (n) of blastocysts are given in each case. Each dot represents data form one. p < 0.05 was accepted as statistically significant

## Discussion

In the present study, we show the mtDNA levels in trophectodermal cells were high in aneuploid blastocysts; however, these mtDNA levels in trophectodermal cells could not predict blastocyst morphology, implantation potential and clinical outcomes.

The ultimate goal for researchers in the field of assisted reproduction is to devise strategies that can improve implantation rates and live birth. Sophisticated morphological grading of blastocyst and pre-implantation genetic screening have helped identify euploid embryos. Even with the elimination of aneuploid embryos, one-quarter of euploid embryos fail to implant, suggesting additional factors may be involved (Seli, 2016; Well, 2017). In recent years, mtDNA levels has gained significant importance in determination of embryo quality and pregnancy outcomes (Cecchino et al., 2019, Humaidan et al., 2018, Kim & Seli, 2019). Mitochondria is involved in various critical cellular processes such as energy generation in the form of ATP, homeostasis, amino acid synthesis and apoptosis. Elevated levels of mtDNA indicate dysfunctional mitochondrial machinery or compensatory mechanism to fulfill the requirements of the ever-growing embryo due to defective organelle (Leese 2002). It is well established that maternal age has detrimental effect on pregnancy outcomes (Miao et al., 2009). Earlier studies on mtDNA levels were low in cumulus cells or the oocytes from in the trophectodermal cells women with advanced age (Boucret et al., 2015; Chan et al., 2005; Onigo et al., 2016). Fragouli and group reported high mtDNA levels in embryos from women with advanced maternal age at the time of oocyte retrieval (Fragouli et.al., 2015; 2017; Ravichandran et al., 2017). However, in this study we did not observe any correlation between maternal age (24-46 years) at the time of oocyte retrieval and mtDNA levels in trophectodermal cells of blastocyst. Similar to our findings, other studies also failed to report any correlation between maternal age with mtDNA levels in the blastocysts (de Los Santos et al., 2018; Diez-Juan et al., 2015; Klimczak et al., 2018; Victor et al., 2017). Together the results indicate, maternal age does not contribute to alterations in mtDNA levels in the blastocysts.

A good quality embryo is vital for the success of ART. Gardner and Schoolcraft grading system is widely used to assess the morphology of the embryo (Gardner et al., 2000). The quiet embryo hypothesis by Leese suggest that good quality embryo have low metabolism, whereas the embryo under stress tend to be more metabolically active (Leese 2002). On the basis of this hypothesis, various studies have analyzed the levels of mtDNA and its association of the developmental competence. Studies have reported high levels of mtDNA in poor quality embryos (de Los Santos et al., 2018; Diez-Juan et al., 2015; Klimczak et al., 2018; Murakoshi et al., 2013). Interestingly, in our study, we did not find any correlation between morphological grading of blastocysts and mtDNA levels in the trophectodermal cells. In concordance with our study, Qui et al., also failed to establish any such correlation (Qui et al., 2018). Thus, association between mtDNA levels in trophectodermal cells and blastocyst quality is debatable. We next tested the levels of mtDNA in trophectodermal cells and its association with ploidy status of the blastocyst. We observed higher levels of mtDNA in trophectodermal cells of aneuploid blastocysts as compared to euploid blastocysts. Interestingly, the high level of mtDNA in trophectodermal cells was restricted to polysomic blastocysts. Our results are consistent with earlier study, wherein polysomic embryos showed elevated levels of mtDNA (Fragouli et.al., 2015). Interestingly, in another study, embryos with monosomies showed higher mtDNA levels as compared to embryos trisomies (de Los Santos et al., 2018). Whereas, some studies fail to show any association between chromosomal status of the blastocyst and mtDNA levels (Qui et al., 2018; Victor et.al., 2017). The association of mtDNA levels and the ploidy status of the blastocyst require further investigation.

Transfer of morphologically and developmentally competent blastocyst to the uterus does not always guarantee pregnancy. Fragouli and group showed euploid embryos with higher mtDNA levels failed to implant, whereas embryos with low mtDNA implanted successfully (Fragouli et al., 2015; 2017). Therefore, we checked if alterations in levels of mtDNA in trophectodermal cells of euploid blastocyst could impact its implantation potential. However, our data failed to establish any correlation between levels of mtDNA in trophectodermal cells and implantation potential of blastocyst. This is in concordance to earlier data where, no correlation such was observed (Qui et al., 2018; Victor et al., 2017). Finally, we asked if the mtDNA levels could predict outcomes of implanted blastocysts. The results revealed the levels of mtDNA in the blastocyst had no correlation with the pregnancy outcomes like miscarriage or live birth. Our study is in concordance to earlier studies where Qui et al., also failed to establish any correlation between mtDNA levels in blastocyst and clinical outcomes (Qui et al., 2018). Taken together, the use of mtDNA levels in the blastocyst as biomarker to predict implantation outcomes and pregnancy remains inconclusive.

From above it is evident that there are differences in reports from different studies. What could be the possible reason for the discrepancies should be addressed. In a recent study, embryos from women with higher body mass index (BMI) showed higher mtDNA copy number; and high levels of maternal serum progesterone inversely correlated with mtDNA levels (de Los Santos et al., 2018). It could be possible that there could be differences in BMI or serum progesterone levels in women in different studies. However, in our study, the baseline BMI and serum progesterone levels did not significantly differ between the different groups and had no effect of mtDNA levels (data not shown). It is possible that technical differences in quantification of mtDNA levels, sample to sample variability, quantifying number of cells during biopsy, sample storage method ethics are some reasons for differences in results (Fragouli et al., 2013; Humaidan et al., 2018; Well, 2017). There is a need of defined selection criteria and standardized procedure to study if there is correlation between mtDNA and ART outcomes.

In summary, the present study fails to support the notion that trophectodermal mtDNA level can predict the embryo quality, implantation ability of the blastocysts or pregnancy outcomes. The limiting factor of our study could be sample size, which fails to account for type II error. Larger study size along with standardized protocol for mtDNA evaluation, quantification and culture methods is the need of hour.

## Acknowledgement

SL is grateful to Council of Scientific & Industrial Research (CSIR), India for Senior Research Fellowship.

## Author’s role

G.V., R.N., M.A., and S.S., the principal investigators take primary responsibility of the paper. G.V., R.N.,M.A., did the clinical work, counselled the patients and collected the samples. RN performed the experiments analyzed the data. D.M., and S.L., contributed in data analysis, data interpretation. All the authors contributed in manuscript preparation.

